# Pathologically distinct fibroblast subsets drive inflammation and tissue damage in arthritis

**DOI:** 10.1101/374330

**Authors:** Adam P Croft, Joana Campos, Kathrin Jansen, Jason D Turner, Jennifer Marshall, Moustafa Attar, Loriane Savary, Harris Perlman, Francesca Barone, Helen M McGettrick, Douglas T Fearon, Kevin Wei, Soumya Raychaudhuri, Ilya Lorsunsky, Michael B Brenner, Mark Coles, Stephen N Sansom, Andrew Filer, Christopher D Buckley

## Abstract

The identification of lymphocyte subsets with non-overlapping effector functions has been pivotal to the development of targeted therapies in immune mediated inflammatory diseases (IMIDs). However it remains unclear whether fibroblast subclasses with non-overlapping functions also exist and are responsible for the wide variety of tissue driven processes observed in IMIDs such as inflammation and damage. Here we identify and describe the biology of distinct subsets of fibroblasts responsible for mediating either inflammation or tissue damage in arthritis. We show that deletion of FAPα+ synovial cells suppressed both inflammation and bone erosions in murine models of resolving and persistent arthritis. Single cell transcriptional analysis identified two distinct fibroblast subsets: FAPα+ THY1+ immune effector fibroblasts located in the synovial sub-lining, and FAPα+ THY1- destructive fibroblasts restricted to the synovial lining. When adoptively transferred into the joint, FAP α+ THY1- fibroblasts selectively mediate bone and cartilage damage with little effect on inflammation whereas transfer of FAP α+ THY1+ fibroblasts resulted in a more severe and persistent inflammatory arthritis, with minimal effect on bone and cartilage. Our findings describing anatomically discrete, functionally distinct fibroblast subsets with non-overlapping functions have important implications for cell based therapies aimed at modulating inflammation and tissue damage.

---

The identification and therapeutic manipulation of leucocyte subsets and the cytokines they produce, has provided the basis for the spectacular success of biologic therapies in immune mediated inflammatory diseases (IMIDs) and cancer^1,2^. However, significant numbers of patients continue to have refractory disease or fail to achieve complete remission^3^. These observations suggest the existence of additional pathogenic pathways, responsible for disease persistence, that have yet to be targeted therapeutically.

Non-hematopoietic, tissue resident stromal cells, contribute to the pathogenesis of many diseases and are known to develop epigenetically imprinted, site and disease specific phenotypes^4-7^. Rheumatoid arthritis (RA) is a prototypic IMID^8^ in which synovial fibroblasts (SFs) play a key role in the destruction of articular cartilage and bone^9,10^ but also modulate the joint microenvironment to favour the persistence of inflammation^11^. However, it remains unknown whether the processes of inflammation and tissue damage are mediated by different subsets of fibroblasts or, instead, reflect cellular plasticity residing within a single fibroblast population^7,13^.

While recent studies have furthered our understanding of the transcriptional heterogeneity of human SFs during inflammation^12,13^, the functional significance of the fibroblast subsets identified in such studies is unknown. A clearer understanding of the biology of SFs is therefore essential to provide a coherent rationale for their therapeutic targeting

We found that expression of fibroblast activation protein α (FAPα), a cell membrane dipeptidyl peptidase^14^, was significantly higher in synovial tissue of patients who fulfilled diagnostic criteria for RA over time, compared to patients in whom joint inflammation eventually resolved (**Fig 1A,B**). Furthermore, expression of the *Fap* gene in cultured SFs isolated from patients with early RA was increased *in vitro* (**Fig 1C**) compared to other diagnostic groups. This suggests that the FAPα+ associated cell phenotype may be pathogenic.

**Fig. 1:**
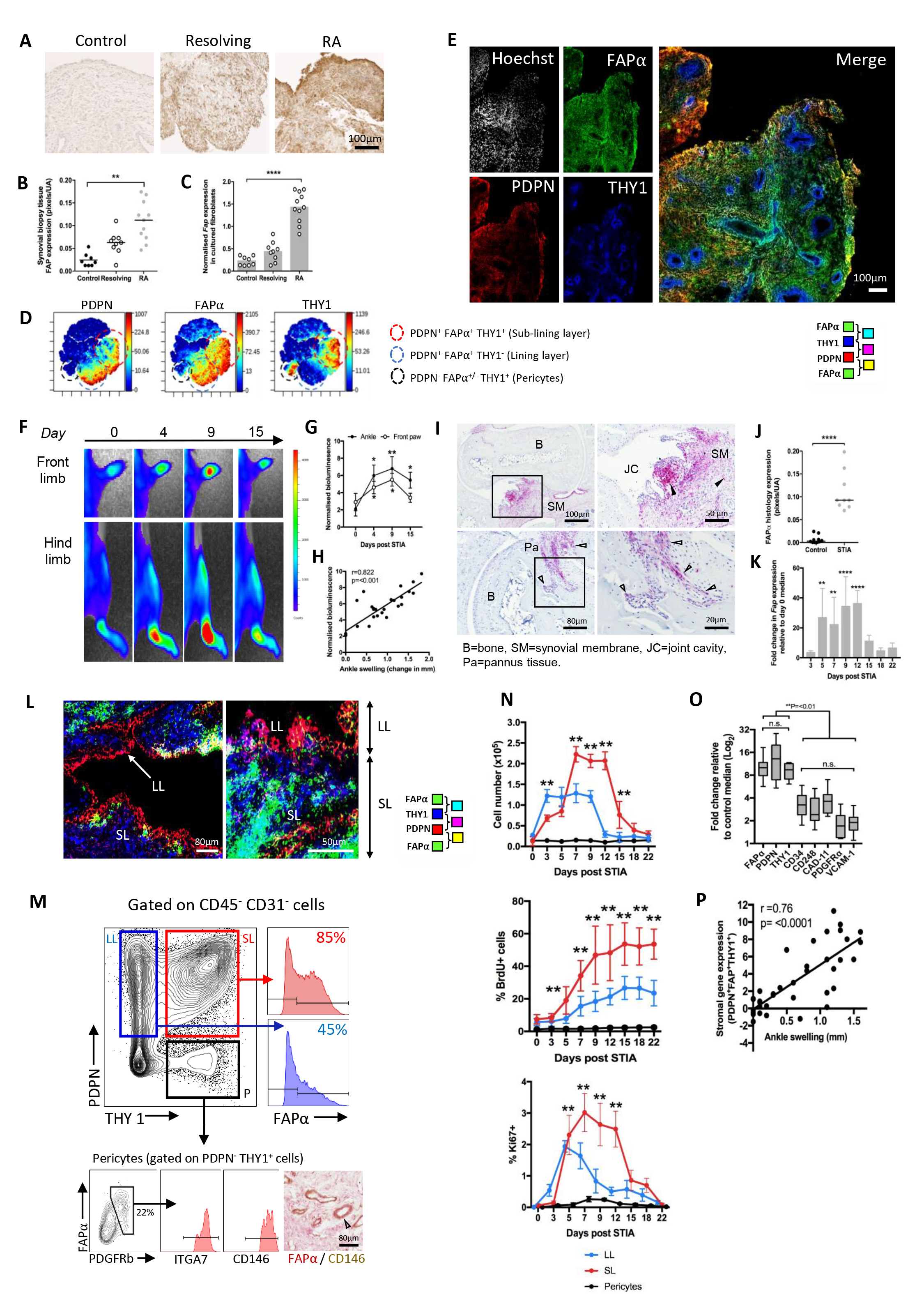
FAPα expressing SFs accumulate within the joint during inflammatory arthritis. **(A)** FAPα expression in synovial biopsy tissue and (**B**) quantification by pixel count (Kruskal-Wallis p=0.0025, Dunn’s post-test). (**C**) Expression of *Fap* transcript by human SFs expanded *in vitro*. 1-way ANOVA and Dunnett’s post hoc compared to control. **(D)** Mass cytometry viSNE plots of RA synovial CD45- cells. (**E**) Confocal microscopy of human RA synovium. (**F**) Bioluminescence in FAPα-luciferase mice during STIA. (**G**) Quantification of bioluminescence signal (n=8, 1-way ANOVA and Dunnett’s post hoc compared to day 0). (**H**) Spearman correlation between bioluminescence and ankle swelling. (**I**) FAPα (red) expression in hind paw joints of STIA mice day 9. (**J**) Quantification of pixel count of FAPα expression, Students t-test. (**K**) Time course analysis of *Fap* transcript in CD45-CD31- SM cells (n=8, 1-way ANOVA and Dunnett’s post hoc compared to day 3). (**L**) Confocal microscopy of day 9 STIA SM. (**M**) Representative flow cytometry plots of LL, SL and pericytes (P) and immunohistochemistry for FAPα and CD146 in day 9 STIA. (**N**) Absolute numbers, percentage of BrdU+ and Ki67+ cells within LL, SL and pericytes (n=6, Students t-test, SL compared to LL). (**O**) Fold change of stromal cell markers, from resting control, by qRT-PCR in SM at day 9 STIA (n=8, 1-way ANOVA and Tukey’s post hoc, ns=not significant). (**P**) Pearson’s correlation between combined expression of *Pdpn*, *Thy1* and *Fap,* and ankle swelling. All data are mean ± SD for at least two independent experiments. ^*^p=<0.05, ^**^p=<0.01, ^***^p=<0.001 and ^****^p=<0.0001.

To map the expression of FAPα in RA synovium we used mass cytometry (CyTOF) analysis of the stromal compartment of the synovial membrane using a combination of podoplanin (PDPN), to identify fibroblasts and THY1 (CD90) to identify sub-lining layer (SL) from lining layer (LL) fibroblasts as validated in previous studies ^12,13,15^. FAPα expression co-localized with PDPN in both the LL and SL cells (**Fig 1D**). A small subset of pericytes (defined as CD45-, PDPN- and THY1+) also expressed FAPα. These findings were confirmed by confocal analysis of RA synovium (**Fig 1E**).

To determine the role of FAPα+ SFs in inflammation, we induced serum transfer inflammatory arthritis (STIA) in a transgenic FAPα luciferase-DTR reporter mouse^16^. FAPα expression increased during the course of arthritis (**Fig 1F,G**). The level of bioluminescence correlated with the severity of joint swelling (**Fig 1H**). Synovial expression of FAPα was either low or undetectable under resting conditions (**extended data 1A**) by histology, but increased throughout the synovial membrane (SM) and focal areas of pannus tissue invading cartilage and bone during inflammation (**Fig 1I,J, extended data 1A**).

During arthritis, FAPα expression was restricted to stromal cells (**extended data 1B,C,D**) and the number of FAPα expressing fibroblasts increased during inflammation returning to baseline levels following resolution (**Fig 1K and extended data 1C,E**), confirming that FAPα is an accurate marker of tissue inflammation (**Fig 1F-K, extended data 1A,C,E)**.

In the murine synovium, THY1 expression also distinguished the SL from LL fibroblasts, with PDPN and FAPα expressed in both cellular compartments (**Fig 1L, M**). To determine how FAPα expressing cell populations changed during arthritis, we quantified their absolute number and assessed their proliferative capacity, by both BrdU incorporation and Ki67 staining using flow cytometry (**Fig 1N**). The severity of joint inflammation positively correlated with an increase in the total number of FAPα+ THY1+ expressing cells but not FAPα+THY1- cells. (**extended data 1F-H**). A significant increase in the proliferation of both THY1- FAPα+ (LL) and THY1+ FAPα+ (SL) cells was observed during inflammation, with THY1+ FAPα+ cells expanding the most (**Fig 1N**). In contrast, there was very little change in the number of FAPα expressing pericytes (**Fig 1N**). *Pdpn*, *Fap* and *Thy1* mRNA showed a significantly higher induction in the inflamed SM compared to resting state (**Fig 1O**) and positively correlated with the severity of ankle swelling (**Fig 1P**). Collectively these data support the idea that expansion of a potentially pathogenic population of SFs marked by a PDPN, FAPα and THY1 expression, occurs in the inflamed synovium.

To determine if FAPα+ fibroblasts play a direct functional role in arthritis we selectively deleted FAPα expressing cells at various stages of arthritis (**extended data Figs 1I-M and 2**). Specific deletion of FAPα expressing cells was confirmed *in vivo* by bioluminescence imaging, histology and flow cytometry (**extended data 1I-M**). The deletion of FAPα expressing cells led to a significant reduction in the cellularity of the synovium with FAPα cell deletion negatively correlating with change in ankle thickness (**extended data 1N,O**).

**Fig. 2.**
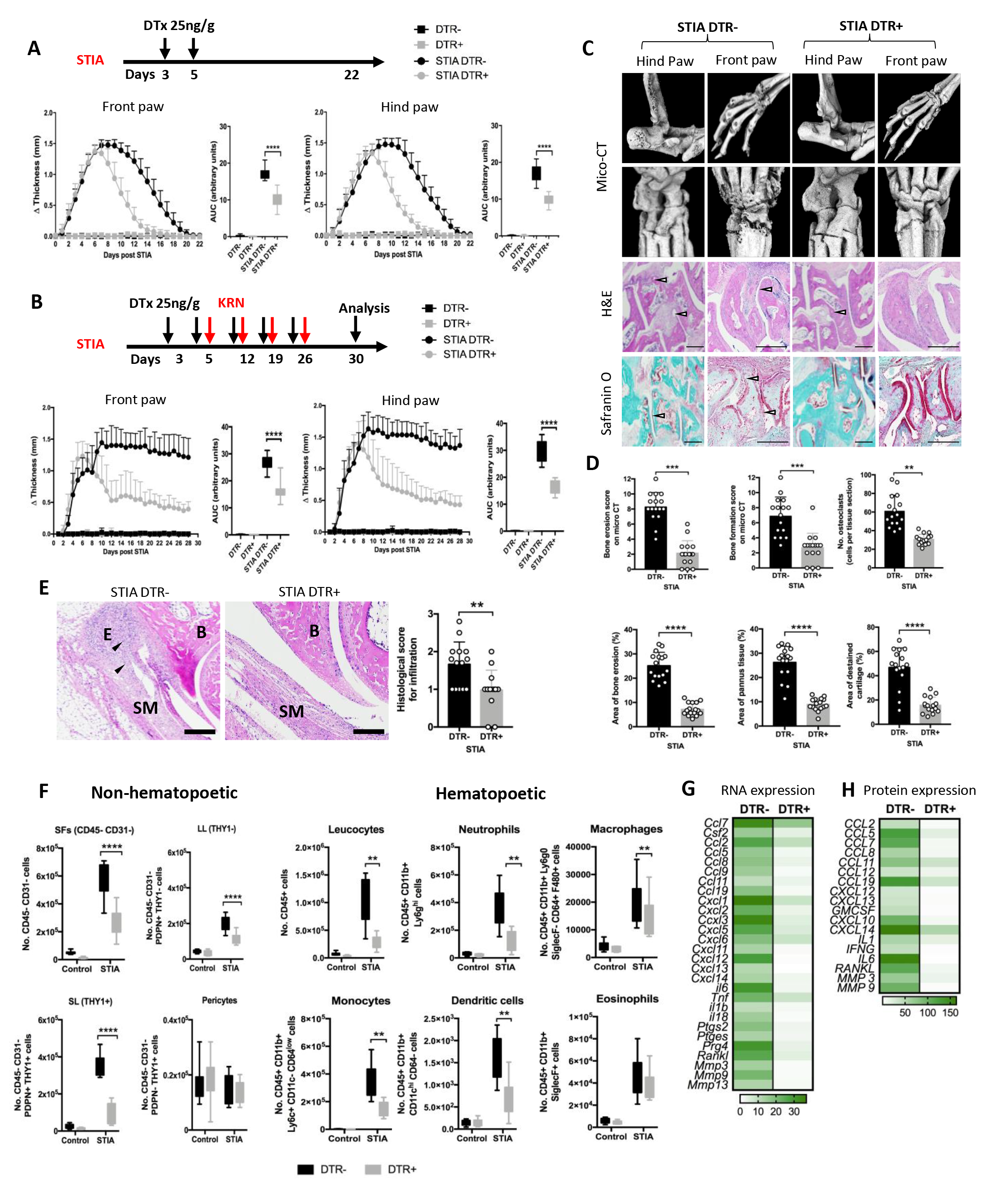
Deletion of FAPα expressing cells attenuates synovial inflammation *in vivo*. (**A**) Synovial inflammation in FAPα deleted (STIA DTR+) versus non-deleted (STIA DTR-) mice during arthritis compared to non-arthritic FAPα deleted (DTR+) and non-deleted (DTR-) mice with area under the curve analysis (AUC). N=6-13, 1-way ANOVA and Dunnett’s post hoc analysis. (**B**) Effect of sustained FAPα cell deletion on synovial inflammation in persistent STIA model compared to non-arthritic mice. N=4- 8, 1-way ANOVA and Dunnett’s post hoc analysis. (**C**) Representative images of day 28 persistent STIA and **(D)** quantification of bone erosion and new bone formation (microCT), osteoclast number per tissue section (example cathepsin K staining, **extended data 4C**), histomorphometric analysis of bone erosion and synovial pannus formation (H&E staining), and cartilage damage (Safranin O depigmentation). Student’s t-test. (**E**) Hind paw H&E staining of day 9 STIA DTR+ and DTR- mice. Arrow shows leucocyte infiltration. SM=synovial membrane, B=bone, E=enthesis. Histological score for leucocyte infiltration. (**F**) Quantification of the number of non-hematopoietic (n=8) and hematopoietic (n=13) cells in digested synovia day 9 STIA analysed by flow cytometry. (**G**) mRNA expression by qRT-PCR in CD45-CD31- cells isolated from DTR- and DTR+ synovial tissue at day 9 STIA (n=8 mice). Expressed as fold change to non-arthritic control mice. (**H**) Luminex analysis in the supernatants of the populations described above, after 24 hours in culture (n=6). Expressed as fold change to non-arthritic control mice. Data are presented as mean ± SD and performed over three independent experiments. ^*^p=<0.05, ^**^p=<0.01, ^***^p=<0.001 and ^****^p=<0.0001. Scale bar=100μm.

Deletion of FAPα expressing cells led to attenuated synovial inflammation and significantly accelerated resolution of inflammation (**Fig 2A).** The same effect was observed regardless of the stage of arthritis at which cells were deleted (**extended data 2A,B**), with the exception of prophylactic deletion of FAPα+ cells prior to induction of arthritis (**extended data 2C**), an observation consistent with the low numbers of FAPα+ cells in the synovial membrane under resting conditions (**extended data 1A**).

The reduction in arthritis following FAPα+ cell deletion was observed in both resolving and persistent models of arthritis (**Fig 2B**). FAPα+ cell deletion resulted in a reduction in structural joint damage (cartilage and bone), leucocyte infiltration and inflammatory bone re-modelling (**Fig 2C,D,E and extended data 3A-C**). Consistent with these findings the number of osteoclasts was also reduced (**Fig 2D, extended data 3D, E**) as was the expression of osteoclast and osteoblast bone markers in digested whole joint tissue (**extended data 3F**). FAPα+ cell deletion was associated with a reduction in both THY1- and THY1+ lining and sub-lining fibroblasts, with no significant change in pericyte numbers (**Fig 2F)**. Circulating blood monocyte number and phenotype were unchanged compared to control mice (**extended data 4A**), excluding any potential indirect effects of myelosuppression following DTx treatment.

Accompanying these changes was a marked reduction in the number of synovial leucocytes, specifically neutrophils, CD11b+ DCs, monocytes and macrophages, but not eosinophils (**extended data Fig 5B** for gating strategy for leucocytes, **Fig 2F, extended data 5C,** persistent model). In addition, deletion of FAPα fibroblasts from the SM resulted in a reduction in the number of macrophages in the LL and a reduction in the percentage of MHC Class II expressing macrophages. Consistent with this observation macrophages from FAPα deleted mice had a more anti-inflammatory phenotype compared to non-deleted mice (**extended Fig 3C-F**). Deletion of FAPα expressing cells was also accompanied by a marked reduction in a panel of pro-inflammatory chemokines and cytokines in FACS sorted CD45- CD31- live cells; both mRNA (**Fig 2G**) and protein (**Fig 2H**). These findings suggest that FAPα expressing cells in the synovium are a significant source of pro-inflammatory cytokines and chemokines, as well as MMPs and RANKL and provides a molecular mechanism for the reduction in clinical inflammation, synovial infiltrate and joint damage observed in the joints of FAPα deleted mice.

We next explored whether both THY1- FAPα+ (LL) and THY1+ FAPα+ (SL) fibroblast populations contribute equally to inflammation and bone damage. We first performed single cell mRNA sequencing of CD45- non-haematopoietic cells from inflamed mouse synovium to obtain an unbiased characterization of the overall transcriptional heterogeneity in SFs at the peak of synovial inflammation. After assigning identities to all cell clusters (**Fig 3A, extended data 5A-D and 6)**, targeted re-analysis of the fibroblast populations only, based on expression of known fibroblast markers, revealed the existence of five phenotypically distinct subgroups **(Fig 3B, extended data 7 and 8A-D).** Gene ontology (GO) analysis of significant cluster marker genes revealed a diversification of function between the subsets: F1 fibroblast marker genes were over-represented in “extracellular matrix organisation and repair”, while F2 and F3 fibroblasts showed an enrichment for the “immune”, “inflammatory response” and “complement activation” biological processes. F4 fibroblasts express genes characteristic of an actively cell cycling population while F5 fibroblasts were over-represented in gene expression associated with “transmembrane transporter activity” **(Fig 3C).** Examination of the top cluster marker genes allowed us to easily differentiate these five subsets at the mRNA expression level (**Fig 3D**). While *Pdpn* and *Fap* were expressed by all of the fibroblast subsets, *Thy1* was expressed selectively by F1-F4 fibroblasts but not F5 fibroblasts **(Fig 3D, extended data 7),** suggesting that we could use THY1 as a marker to discriminate the lining layer F5 subset from the four sub-lining subsets (F1-F4). We also examined the expression of other known fibroblast markers across the subsets as well as the specific expression of selected chemokines (**extended data 8A,C**).

**Fig. 3.**
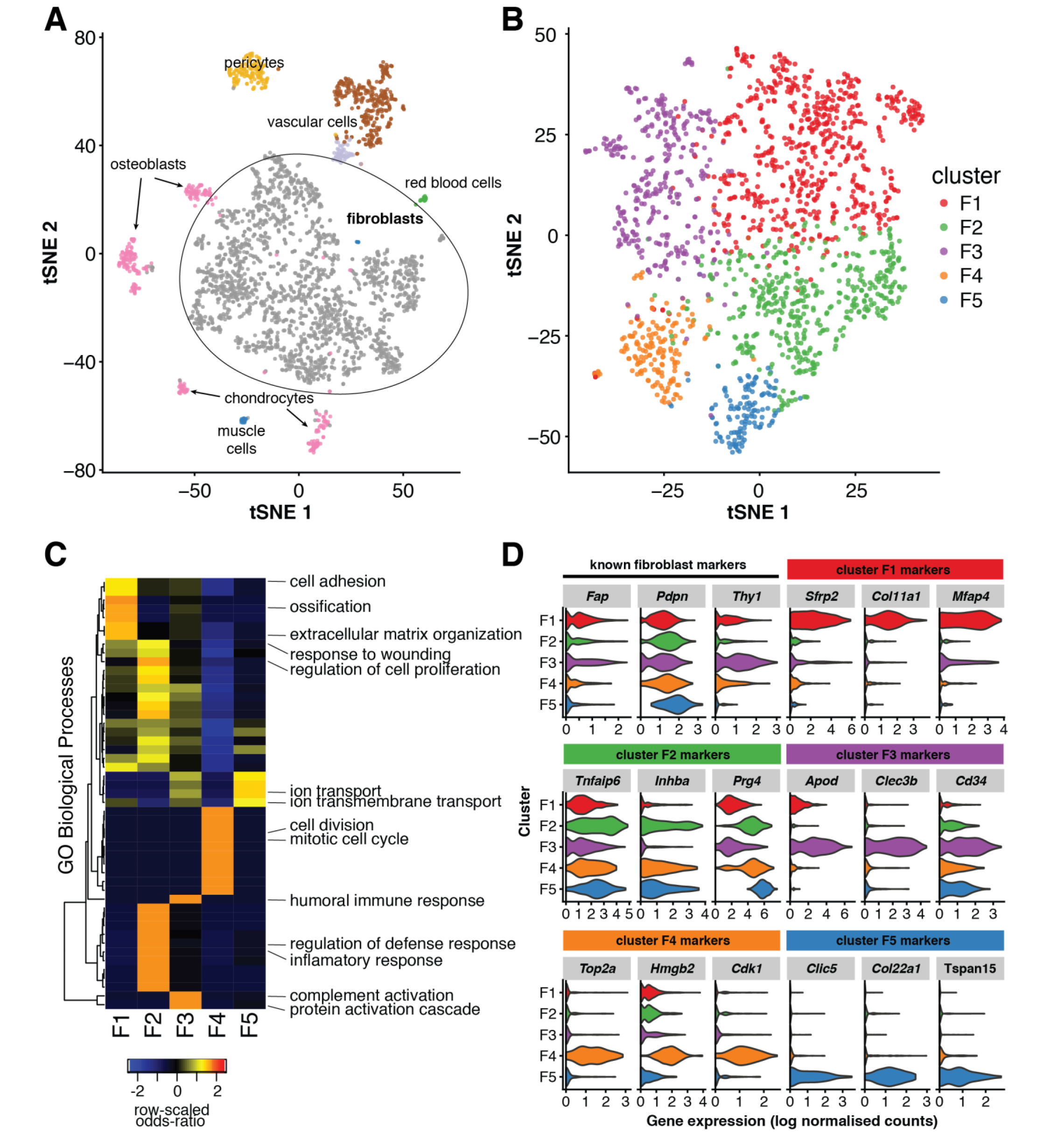
Single-cell RNA-sequencing reveals distinct fibroblast subsets with specific transcriptional profiles. **(A)** The t-SNE projection shows 2814 non-hematopoietic (CD45^-ve^ CD31^-ve^ cells) stromal cells FACS sorted from inflamed synovium digests from day 9 STIA mice (n=3 biological replicates). Group of cells are coloured according to major cell type as identified on the basis of the expression of known marker genes (see **Extended data 6 and 7**). (**B**) Re-analysis of the fibroblast population (n=1725 cells) reveals five distinct subsets in the inflamed STIA joint, here denoted F1-F5. Unsupervised graph-based clustering was performed with Seurat^22^. (**C**) Over-representation analysis of Gene Ontology (GO) categories suggests that the five subsets may have different biological functions. Each row of the heatmap represents a biological process that was significantly over-represented in at least one cluster (p < 0.05, one-sided Fisher’s exact tests). (**D**) Expression of marker genes in the identified fibroblast clusters. The first three panels show expression of known pan-fibroblast markers. The remaining panels show genes discovered to be conserved, significant marker genes for each fibroblast cluster F1-F5 (adjusted p value < 0.05 in each of the three replicates, Wilcoxon tests).

To confirm the validity of using PDPN, THY1 and FAPα as a cassette of cell surface markers able to discriminate LL and SL, we performed ultra-low input RNA sequencing on sorted PDPN+, FAPα+/- and THY1+/- cell populations (**extended data 9A-C, 10**). Principal components analysis (PCA) of transcriptional differences confirmed that the subsets defined by expression of THY1 represented transcriptionally distinct populations (**extended data 9A,** gating strategy for sort purified cell populations). The most obvious separation was between THY1+ versus THY1- populations (regardless of FAPα expression (**extended data 9B**). The THY1+ cell population showed expression of many chemokines and cytokines and expressed F1-F4 subset specific genes (**extended data 9C and extended data 10**). In contrast, THY1- cell gene expression was consistent with F5 fibroblasts associated genes such as *Prg4*, *Clic5* and *Tspan15* as well as genes associated with cartilage and bone erosion (**extended data 9C and extended data 10**). Therefore the greatest differences in the transcriptional profiles of SFs is based on expression of THY1 representing their location in either the lining or sub lining of the synovial membrane

As predicted from the single cell transcriptome analysis, FAPα+ THY1+ subsets expressed an immune effector profile with higher expression of chemokines as well as cytokines including: *Il-6*, *Lif*, *Il33* and *Il34* **(extended data 10)**. In contrast, FAPα+THY1- expressing SFs expressed higher levels of *Ccl9* and RANKL,, both potent inducers of osteoclast activity, as well as *Mmp3*, *Mmp9* and *Mmp13*; matrix metalloproteinases involved in cartilage degradation (**extended data 10**). These findings were validated for protein expression using Luminex analysis of the culture supernatants of *ex vivo* purified stimulated FAPα expressing THY1+ and THY1- cells (**Fig 4A**). Moreover, FAPα+ THY1- cells expressed RANKL on their cell surface, secreted higher levels of RANKL and exhibit a significantly increased RANKL/OPG ration compared to FAPα+ THY1+ (**Fig 4B,C)**. Taken together these results strongly suggested that the THY1- and THY1+ cells might perform distinct non-overlapping functions *in vivo.*

**Fig. 4.**
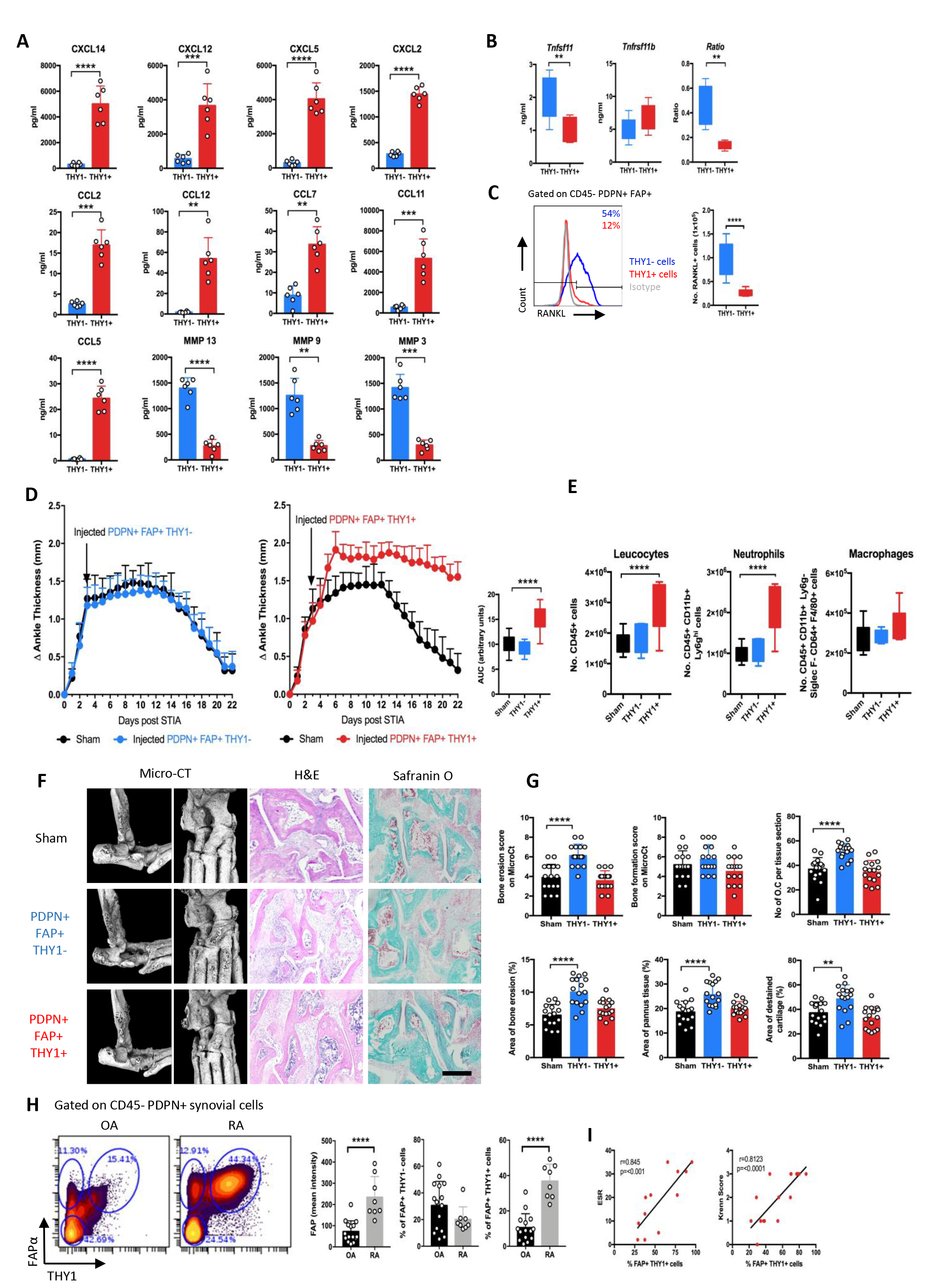
Distinct fibroblast subsets are responsible for different aspects of disease pathology in joint inflammation. (**A**) Luminex analysis of supernatants from IL-1β stimulated FAPα+THY1- (LL) and FAPα+THY1+ (SL) cells isolated from day 9 STIA synovia (n=6), including (**B**) secretion of RANKL (*Tnfsf11*) and OPG (*Tnfrsf11b*) (n=6). (**C**) Cell surface expression of RANKL by flow cytometry and absolute numbers of RANKL+ THY1-/+ cells in inflamed synovia (n=8). (**D**) Effect on ankle swelling of adoptive transfer of 500,000 sort purified PDPN+FAPα+THY1- (blue) and PDPN+FAPα+THY1+ (red) cells, by intra-articular injection into inflamed ankle joints at day 3 STIA (n=8). (**E**) Flow cytometric analysis of absolute number of hematopoietic compartment in digested synovial tissue from injected joints at day 12 (n=8, 1-way ANOVA with Dunnett’s post hoc compared to sham control). (**F**) Representative images at day 12 and **(G)** quantification of bone erosion and new bone formation (microCT), osteoclast number (cathepsin K staining), bone erosion and synovial pannus formation (H&E staining), and cartilage damage (Safranin O depigmentation). (n=8, 1-way ANOVA and Dunnett’s post hoc compared to sham). (**H**) Flow cytometric analysis of PDPN+ FAPα+ and THY1+ in synovium of OA and RA patients (gated on CD45- CD31- cells). Mean fluorescence intensity for FAPα expression, percentage of each fibroblast subset within the synovia. (**I**) Spearman correlation between THY1+ FAPα+ cells in RA and serum ESR level and synovial Krenn score. All data are mean ± SD for at least two independent experiments. ^*^p=<0.05, ^**^p=<0.01, ^***^p=<0.001 and ^****^p=<0.0001. Scale bar=100μm.

To directly test this hypothesis we adoptively transferred PDPN+ FAPα+ THY1- or PDPN+ FAPα+ THY1+ cells directly into the inflamed ankle joint of mice. Injection of PDPN+ FAPα+ THY1+ cells resulted in more severe and sustained paw swelling (**Fig 4D**), with higher levels of leucocyte infiltration compared to sham injected joints (**Fig 4E**) but with little effect on bone and cartilage destruction **(Fig 4F,G)**. In contrast, the injection of PDPN+ FAPα+ THY1-cells had no effect on the severity or temporal dynamics of joint inflammation **(Fig 4D),** but did result in increased osteoclast activity, increased bone erosion and structural joint damage (**Fig 4F,G**). Collectively, these data suggest that in pathological conditions PDPN+ FAPα+ THY1+ expressing cells assume an immune effector role capable of sustaining inflammation through the production of a distinct repertoire of chemokines and cytokines, whereas PDPN+ FAPα+ THY1-cells are bone effector cells that mediate joint damage.

In support of this conclusion, and as validation of the relevance of our findings to human disease, we identified an expanded population of PDPN+ FAPα+ THY1+ immune effector fibroblasts in the synovia of patients with RA, in whom joints are persistently inflamed, compared to patients with osteoarthritis (OA), a disease characterised predominately by joint damage rather than inflammation (**Fig 4H**). The expansion of THY1+ FAPα+ cells positively correlated with markers of systemic inflammation including circulating Erythrocyte Sedimentation Rate (ESR) and the tissue Krenn score, a histological measurement of synovial inflammation^17^ (**Fig 4I**).

In conclusion we have identified and described the pathological significance of fibroblast heterogeneity in RA, an IMID in which inflammation and damage play key pathogenic roles. We describe discrete, anatomically distinct subsets of fibroblasts with non-overlapping effector cell functions including joint damage (production of MMPs and induction of osteoclastogenesis) and immuno-inflammatory regulation (production of inflammatory cytokines and chemokines). These findings provide the unpinning justification for the development of therapies that selectively target deletion or replacement of different mesenchymal subpopulations in a wide range of diseases.

**Supplementary Information** is linked to the online version of the paper at

## Acknowledgements

A.P.C was supported by a Wellcome Trust clinical career development fellowship #WT104551MA; A.F by Arthritis Research UK clinician scientist fellowship #18547; F.B by an Arthritis Research UK senior fellowship; K.W. by Rheumatology Research Foundation Scientist Development Award; K.J. by a Wellcome Trust PhD studentship; S.N.S. and M.A. are supported by the Kennedy Trust for Rheumatology Research. This work was supported by the Arthritis Research UK Rheumatoid Arthritis Pathogenesis Centre of Excellence #20298; The National Institutes of Health Accelerating Medicines Partnership in RA/SLE and Arthritis Research UK programme grant #19791 (to C.D.B). This paper presents independent research supported by the NIHR Birmingham Biomedical Research Centre at the University Hospitals Birmingham NHS Foundation Trust and the University of Birmingham. The views expressed are those of the author(s) and not necessarily those of the NHS, the NIHR or the Department of Health.

## Author Contributions

A.P.C. conceived the project, performed experiments, analysed data, and wrote the manuscript. J.C. performed experiments, analysed data, and wrote the manuscript. J.M, L.S, J.D.T, K.W, H.M.M. and I.L. performed experiments and analysed data. K.J analysed the single cell RNA sequencing data and helped to write the manuscript. M.A. performed single cell capture and library preparation.. KW analysed mass cytometry data. AF participated in study design, patient recruitment, sample acquisition, and review of the data. S.N.S supervised the design, execution, analysis and interpretation of the single-cell transcriptomics experiment and helped write the manuscript. F.B. H.P., D.T.F, S.R, M.B. and M.C. provided critical interpretation of the data and contributed to writing the manuscript. A.F. and C.D.B. conceived the project, supervised the work, analysed data, and co-wrote the manuscript. All authors discussed the results and commented on the manuscript.

## Author Information

RNA sequencing data is available at the GEO repository, accession number xxx. The authors have no competing financial interests. Correspondence and requests for materials should be addressed to C.D.B.

## METHODS

### Human patient cohorts

The study protocols are Institutional Review Board approved at Partners HealthCare, Hospital for Special Surgery, and the University of Birmingham Local Ethical Review Committee via approved protocols with appropriate informed consent, as required. Synovial tissues and clinical outcome data of patients included in the early arthritis patient cohort in Birmingham (BEACON) were used in this study. Patients with clinical synovitis present in at least one synovial joint (of 66 joints examined) were recruited to the BEACON cohort if symptom duration, defined as any symptom attributed by the assessing rheumatologist to inflammatory joint disease (pain, stiffness and/or swelling), was no greater than 3 months. All patients were naïve to treatment with disease-modifying anti-rheumatic drugs (DMARDs) and corticosteroids at inclusion. Diagnostic and prognostic outcomes were assigned after 18-months of follow-up. Patients were classified as having RA according to the 2010 ACR/EULAR classification criteria for RA^18^ or were classified as having self-limiting resolving disease if they had no clinical evidence of synovial swelling and had not taken DMARDs or received glucocorticoid treatment in any form in the preceding 3 months. Healthy (control) synovial tissue was obtained from uninflamed joints of patients with joint pain and normal imaging studies who were undergoing exploratory arthroscopy. OA samples were obtained by synovial biopsy of affected joints.

### Human synovial tissue analysis

Synovial tissue was obtained by ultrasound guided synovial biopsy of the inflamed joint and tissue was collected and processed as described below. Synovial tissue samples were snap-frozen in Tissue-Tek OCT medium (Miles, Elkhart, IN) immediately after collection. In order to account for heterogeneity, six to eight biopsies from different areas of the joint were combined in one block of tissue. Cryostat sections (5μm) were cut, mounted on Star Frost adhesive glass slides (Knittelplaser, Baunschweig, Germany) and stored at -80°C.

For immunohistochemical staining, acetone fixed slides were re-hydrated in PBS (Sigma-Aldrich) for 10 minutes prior to blocking with Bloxall reagent (Vector Laboratories, California USA) for ten minutes, followed by 10% normal horse serum (Sigma-Aldrich) in PBS for a further 10 minutes. Slides were stained using sheep anti-FAP antibody (R&D, AF3715) for 1 hour at room temperature (RT). Slides were washed before the application of donkey anti goat HRP (Dako). Horseradish peroxidase (HRP) staining was developed using the ImmPACT DAB Peroxidase HRP Substrate (Vector Labs) and counterstained with haematoxylin. Slides were mounted in VectaMount (Vector Labs) before imaging using the Zeiss Axio Scan and analysis using Zen lite 2012 software (ZEISS).

For immunofluorescence staining acetone fixed slides were rehydrated in PBS and blocked with 10% normal human serum for 10 min at RT. Incubation with primary antibodies was for 1 hour at RT. As primary antibodies, anti-FAP antibody (mouse IgG1, clone F11-24; eBioscience), anti-PDPN (Rat IgG2a, clone NZ-1.3 eBioscience or mouse IgG1, clone D2-40; AbD Serotec, Kidlington, UK) and anti-THY1 (F15-42-1, Merck Millipore, and clone Thy-1A1, R&D Systems) were used.

### Human fibroblast cell culture

Human fibroblasts were isolated as previously described^19^ and cultured in Dulbecco’s modified Eagle’s medium (DMEM) (Sigma-Aldrich) supplemented with 2% fetal bovine serum (FBS; Gemini), 2 mM L-glutamine, antibiotics (penicillin and streptomycin), and essential and non-essential amino acids (Life Technologies). The cells were expanded for 3- 20 days for assays in vitro. Fibroblast lines at passage 3 or 4 were used for *in vitro* experiments.

### Mass cytometry

Cryopreserved disaggregated human synovial cells were thawed into RPMI + 10% FBS (HyClone). Viability was assessed with cisplatin (Fluidigm) for synovial cells. Cells were then washed and stained with primary antibody cocktails at 1:100 dilution (CD45, metal 89Y, clone HI30; PDPN, metal 156Gd, clone NC-08; FAP, metal 147Sm, Poly; THY1, metal 162Dy, clone 5E 10). All antibodies were obtained from the Longwood Medical Area CyTOF Antibody Resource Core (Boston, MA). Cells were then washed, fixed and permeabilized using the Ebioscience Transcription Factor Fix/Perm Buffer for 45 minutes, washed in PBS/1%BSA (Sigma-Aldrich)/0.3% saponin, then stained for intracellular markers. Cells were re-fixed in formalin (Sigma-Aldrich), washed with Milli-Q water, and analyzed on a Helios (Fluidigm) for synovial cells. Mass cytometry data were normalized using EQ^™^ Four Element Calibration Beads (Fluidigm) viSNE analyses were performed on cytometry data from 3 of 6 synovial tissue samples, using the Barnes-Hut SNE implementation on Cytobank (www.cytobank.org). All 3 individual synovial tissue sample analyses are shown.

### Mice

All experiments were performed under guidelines of, and were approved by, the UK Home Office and the animal ethics committee of the University of Birmingham. All C57BL/6 mice were purchased from the Jackson Laboratory and bred and housed at a barrier and specific pathogen-free facility at the Biomedical Services Unit, University of Birmingham (Birmingham, UK). Eight to ten week old mice were used for all experiments.

FAP-DTR transgenic (Tg) embryos were a kind gift from Prof Douglas T Fearon. FAP-DTR mice were generated to express the Diphtheria Toxin Receptor (DTR), luciferase reporter (Luc2) and mCherry under the murine fap promoter, by insertion of a 206kb BAC clone (RP23-16A15)^16^. FAP-DTR+ males were bred with albino C57BL/6 females (C57BL/6J-*Tyr^c-2J^*) and heterozygous animals were used for experiments. For FAP+ cell deletion, Diphtheria Toxin (DTx) (List Biological Laboratories) was administered by i.p. injection to 8 week old FAP-DTR mice at a dose of 25ng/g, twice a day, in a prophylactic regime (at days -7 and -5 post STIA) and in a therapeutic regime (at days 3 and 5 or at days 5 and 7 post STIA or at days 10 and 12 post STIA). Sterile water for injections was used as vehicle control.

Serum transfer inflammatory arthritis (STIA) was induced by intravenous injection of 100μl of arthritogenic serum from 8-week-old progeny of KRN mice (K/BxN) mice20. Change in ankle or paw thickness was monitored using calipers (Fowler Tools of Canada), and was reported as ankle/paw thickening: the change in ankle/paw thickness from day 0. Disease severity was quantified using the area under the curve of ankle/paw thickening calculated using Prism 7 software.

In addition, clinical score was assessed daily as a sum of the clinical score for each paw (0 – no arthritis; 1 – mild arthritis, foot maintains V-shape; 2 – moderate arthritis, foot no longer maintains V-shape; 3 – severe arthritis). Each experiment was performed 2–5 times to confirm reproducibility. Whenever possible scoring was performed in a blinded manner by one observer. In the persistent model of arthritis, mice were administered arthritogenic serum once a week and DTx injections twice a week, for a period of 4 weeks. For the purpose of BrdU incorporation, mice were injected at day 0 with 100μl of 10mg/ml BrdU in PBS and then kept on BrdU-containing drinking water (0.8mg/ml).

### In vivo imaging

Mice were intraperitoneally injected with 150 μg/g body weight D-luciferin (PerkinElmer) and imaged using IVIS (Xenogen). Whole body images were acquired for 20 s at 1.5-cm stage height. Data were analysed, normalised and quantified using Living Image (PerkinElmer).

### Flow cytometry and cell sorting

For flow cytometric analysis, murine limbs were dissected and collected in 2ml of RPMI-1640 (+2% FCS) on ice. Muscle and soft tissues were dissected leaving bones and joints intact, these were then transferred into 1.5ml of Digestion Buffer 1 – RPMI-1640 (Sigma-Aldrich) (+2% FCS) containing 0.1g/ml Collagenase D (Roche) and 0.01g/ml of DNase I (Sigma-Aldrich). Samples were incubated at 37°C in a water bath for 40 minutes with stirring and centrifuged at 300g for 4 minutes. Fresh Digestion Buffer 2, 1.5ml RPMI-1640 (+2% FCS) containing 0.1g/ml Collagenase Dispase (Roche) and 0.01g/ml DNase I, was added to the samples and contents were gently mixed and incubated at 37°C for 20 minutes with magnetic stirring. Finally, 15μl of 0.5M EDTA (Sigma Aldrich) was added to the samples followed by incubation at 37°C for 5 minutes. Cell suspensions were filtered through 70μm cell strainers into a 50ml falcon tube containing RPMI-1640 (+2% FCS) and resulting contents centrifuged at 300g for 4 minutes at 4°C. Cell pellets were suspended in PBS containing 0.25g BSA and 0.5M EDTA, for subsequent staining. Cells were stained at 4°C, fixed with BD Cytofix/Cytoperm^™^ and samples acquired with a BD LSR Fortessa. Dead cells were excluded using Zombie Yellow staining (77168, BioLegend). Antibodies used were anti-CD45 (30-F11, ebioscience), anti-Thy1 (53-2.1, ebioscience), anti-podoplanin (8.1.1, ebioscience), anti-FAP (AF3175, R&D systems), mouse anti-goat/sheep IgG biotin (GT-34, Sigma Aldrich), anti-CD31 (390, ebioscience), anti-CD11b (M1/70, ebioscience), anti-Ly6G (1A8, BioLegend), anti-SiglecF (1RNM44N, ebioscience), anti-CD64 (X54-5/7.1, BioLegend), anti-F4/80 (BM8, BioLegend), anti-CD11c (N418, ebioscience), anti-RANKL (IK22/5, BioLegend), anti-BrdU (3D4, BD Biosciences), anti-Ki67 (11F6, BioLegend), anti-CD140b (APB5, BioLegend), anti-ITGA7 (334908, ebioscience), anti-CD146 (ME-9F1, BioLegend), anti-CD115 (AFS98, Bioledgend), anti-CD43 (S11, Bioledgend), anti-MHC Class II (39-10-8, Bioledgend) and anti-Ly6C (HK1.4, Bioledgend). BrdU staining was performed according to manufacturer’s instructions using a BrdU Flow Kit (BD Pharmigen^™^). For cell sorting, a MoFlo Astrios EQ machine (Beckman Coulter) was used immediately after flow cytometry staining.

### Generation and analysis of droplet-based single cell RNA sequencing data

Following sorting, digested, CD45^-ve^ live synovial cells isolated from hind limbs of day 12 STIA inflamed mouse joints (n=3 biological replicate samples, each comprised of cells from the joints of three animals) were captured with the 10X Genomics Chromium system. Sequencing libraries were generated using the 10x Genomics Single Cell 3’ Solution (version 2) kit and subjected to Illumina sequencing (HiSeq 4000, read 2 sequenced to 75bp). Alignment, quantitation and aggregation of sample count matrices was performed using the 10x Genomics Cell Ranger pipeline (version 2.1.0) and mouse reference sequences (version 2.1.0), retaining a median of 59.9k reads/cell (mapped-read depth normalization applied). To circumvent known index-hopping issues with the HiSeq 4000 platform^21^ cell barcodes common to more than one sample were removed from the aggregated count matrix. The UMI count matrix was also randomly down-sampled to ensure normalization of the median number of per-cell counts between the samples. Downstream analysis was performed using the Seurat R package (version 2.3.0) ^22^Butler:2018ex} as follows. Cells with greater than 5% mitochondrial reads or fewer than 500 genes were excluded from the analysis. Cells were down-sampled to a common number: for the full analysis we retained n=938 cells per replicate, while n=575 cells were retained per replicate for re-analysis of the fibroblasts. Per-cell counts were normalised, scaled and the effects of total UMI counts and percentage of mitochondrial counts regressed out. For the fibroblast re-analysis, the difference between G2M and S phase was also regressed out based on the expression of known cell cycle marker genes^23^. In both cases, we retained 30 principle components for tSNE projection and clustering analysis (original Louvain algorithm, resolution set to 0.6 for the full analysis and to 0.4 for the fibroblast reanalysis). Conserved cluster markers were identified as the intersection of those that were significant in separate tests of the cells from the each replicate (Wilcoxon test, BH adjusted p values < 0.05). Only genes found in 10% of cells (either within or outside the cluster of interest) and that showed a minimum log fold difference of 0.25 were tested for differential expression. Geneset over-representation analysis of cluster marker genes was performed using one-sided Fisher’s exact tests (as implemented in the “gsfisher” R package https://github.com/sansomlab/gsfisher) with Biological Process gene sets obtained from the Gene Ontology (GO) database^24^. For this analysis cluster-specific gene universes were defined as those genes expressed in 10% percent of cells (either within or outside the cluster of interest). The computational analyses were performed using the “pipeline_cellranger.py” and “pipeline_seurat.py” pipelines (https://www.github.com/sansomlab/tenx/).

### Bulk RNA sequencing

#### RNA-seq library preparation and sequencing

cDNA synthesis was performed on isolated RNA using the SMART-Seq^®^ v4 Ultra^®^ Low Input RNA Kit for Sequencing. Libraries were pooled and sequenced paired end 75 bases each end with the NextSeq 500 system.

#### RNA-seq gene expression quantification

The reads were mapped to the GRCm38 (Ensemble release 85) mouse genome using STAR alignment software version v2.5.2b^25^. Read counts per gene were produced by the same software. Sample normalization and differential expression analysis was performed using DESeq2 R Bioconductor package^26^. log2 values of read counts regularised by DESeq2 were used in heatmaps in extended data 11 and 12.

#### Luminex analysis

Sort purified PDPN+ FAPα+ THY1- or PDPN+ FAPα+ THY1+ cells were stimulated with 1ng/ml recombinant mouse IL-1β (Peprotech) *ex-vivo* in culture medium for 1 hour. Fresh culture media was then applied and the subsequent supernatant was harvested after 12 hours and analysed using custom selected multiplex bead based assays (Luminex assay panel, R&D systems).

#### Blood biochemistry

Murine blood was collected under terminal anaesthesia by cardiac puncture into heparinized tubes. Plasma was collected and analysed using the APX Pentra 60 (Horiba) according to the manufacturer’s instructions.

#### RNA extraction, cDNA synthesis, and qPCR

RNA was isolated from single cell suspensions of digested synovia, flow sorted or cultured cells using the PicoPure RNA isolation kit (Thermo Fisher Scientific) according to the manufacturer’s instructions. For tissue analysis, wrist or ankle joints were snap frozen in liquid nitrogen at the time of dissection and stored at -80°C. Joints were pulverised to a fine powder in liquid nitrogen using a FreezerMill 6770 (Spex Sample Prep). mRNA was extracted from tissue using ReliaPrep^™^ RNA Tissue Miniprep System (Promega). cDNA synthesis was performed on all samples using iScript cDNA synthesis kit (Bio-Rad) and carried out on a Techne 312 thermal cycler PCR machine.

In order to amplify small amounts of cDNA without introducing amplification bias, TaqMan PreAmp Master Mix was used according to manufacturer’s instructions for flow sorted cell samples. Samples were pre-amplified using 2.5μl of 1:100 diluted pool of primers and probes, 2.5μl of cDNA sample and 5μl of TaqMan^™^ PreAmp Master Mix in a final volume of 10μl. Cycling conditions were as follows: 10 min of enzyme activation at 95°C, 10 cycles of denaturing at 95°C for 15 s plus annealing at 60°C for 4 min, and 10 min of enzyme inactivation at 99°C.

Quantitative RT−qPCR was performed using Taqman assays and Taqman universal Mastermix (both from Applied Biosystems) on a real-time PCR detection system (CFX96 Touch™ Real-Time PCR Detection System). All expression levels were normalized to an internal housekeeping (HK) gene (RPLP0 for human samples and Hprt for mouse samples) and calculated as 2^−(CTHK−CTgene). TaqMan primer/probes (Applied Biosystems) used were Fapα (human: Hs00990791_m1; mouse: Mm01329177_m1), Pdpn (Mm01348912_g1), Thy1 (Mm00493681_m1), CD34 (Mm00519283_m1), CD248 (Mm00547485_s1), Cdh11 (Mm00515466_m1), Pdgfra (MM00440701_m1), Vcam1 (Mm01320970_m1), Ccl7 (Mm00443113_m1), Csf2 (Mm01290062_m1), Ccl2 (Mm00441242_m1), Ccl5 (Mm01302427_m1), Ccl8 (Mm01297183_m1), Ccl9 (Mm00441260_m1), Ccl11 (Mm00441238_m1), Ccl19 (Mm00839967_g1), Cxcl1 (Mm04207460_m1), Cxcl2 (Mm00436450_m1), Cxcl3 (Mm01701838_m1), Cxcl5 (Mm00436451_g1), Cxcl6 (Mm01302419_m1), Cxcl11 (Mm00444662_m1), Cxcl12 (Mm00445553_m1), Cxcl13 (Mm04214185_s1), Cxcl14 (Mm00444699_m1), il18 (Mm00434226_m1), Ptgs2 (Mm00478374_m1), Ptges (Mm00452105_m1), Prg4 (Mm01284582_m1), Rankl (Mm00441906_m1), Mmp3 (Mm00440295_m1), Mmp9 (Mm00442991_m1), Mmp13 (Mm00439491_m1), Ctsk (Mm00484039_m1), Runx2 (Mm00501584_m1), Spp1 (Mm00436767_m1), Acp5 (Mm00475698_m1), Tnfrsf11a (Mm00437132_m1), Sost (Mm00470479_m1), BGLAP (Mm03413826_mH), Dmp1 (Mm01208363_m1), il6 (Mm00446190_m1), Tnf (Mm00443258_m1), il1b (Mm00434228_m1), il10 (Mm01288386_m1), inos (Mm00440502_m1) and Arg1 (Mm00475988_m1).

#### Histology

Murine freshly dissected limbs were either formalin fixed and paraffin embedded (FFPE) or were placed in a cryomold, embedded in OCT compound (both from Sakura Finetek), frozen in dry ice and stored at - 80°C. Frozen sections were then obtained with a tape transfer system CryoJane^®^ (Leica Biosystems), according to manufacturer’s instructions. Immunohistochemistry staining for PDPN (hamster anti-mouse PDPN, ebioscience eBio8.1.1 (8.1.1), FAPα (R&D, AF3715), F4/80 (rat anti-mouse F4/80, BioRad CI:A3-1 and cathepsin K (Abcam, rabbit polyclonal) was performed on FFPE sections, as described above. H&E staining was performed at the Royal Orthopaedic Hospital Pathology Laboratories according to standard protocol. Safranin O staining for cartilage was performed as previously described^27^. Pixel counts were performed using Image J and colour pixel counter plugin. Immunofluorescence staining for podoplanin, FAP (R&D, AF3715) and Thy1 was performed in frozen sections, as described above. Tissue sections were imaged using the AxioScanZ.1 slide scanner.

#### Histomorphometry scoring

Histomorphometric analysis of H&E and Safranin O stained tissue was performed using Zen image analysis software, as previously described 28. Osteoclast were detected by both morphology and cathepsin-K staining on tissue sections and following imaging of whole tissue sections the number of osteoclasts were counted.

#### Adoptive transfer of SF

Adoptive transfer of sort purified cells was performed immediately post sorting by intra-articular injection of 500,000 live cells into inflamed ankle joint (day 4 STIA). Cells were suspended in sterile PBS at a concentration 500,000 cells/50μl. The same volume of PBS was injected in the contralateral joint as a sham control.

#### Micro-CT

Hind limbs were imaged using a Skyscan 1172 microCT scanner (Bruker) using settings and reconstruction algorithms using MeshLab v1.3.2 (open source software developed with the support of the 3D-CoForm project) as previously described^27^. Micro-CT meshes were visualised in MeshLab and meshes were divided into 3 regions: Heel (compromising the calcaneus, centrale, distal tarsals, tibulae and astagalus and distal tibia and fibula), metatarsals and phalanges (excluding the claws). Each region was scored for erosion (0 = normal, 1 = roughness, 2 = pitting, 3= full thickness holes) and the extent of the area affected (0 = none, 1 = a few small areas, 2 = multiple small-medium areas, 3 = multiple medium-large areas). The 2 scores were then multiplied together for each region to give a maximum score per paw of 27. The average score and standard deviation from the 2 blinded scorers are shown.

#### Statistical analysis

Statistical analysis was performed as described in each section using Prism 7 software. Data is presented as Mean ± SD from data obtained from at least two independent experiments.

## REFERENCES

1. Baker, K. F. & Isaacs, J. D. Novel therapies for immune-mediated inflammatory diseases: What can we learn from their use in rheumatoid arthritis, spondyloarthritis, systemic lupus erythematosus, psoriasis, Crohn’s disease and ulcerative colitis? Ann. Rheum. Dis. 77, 175–187 (2018).

2. Smolen, J. S. & Aletaha, D. Rheumatoid arthritis therapy reappraisal: strategies, opportunities and challenges. Nat Rev Rheumatol 11, 276–289 (2015).

3. Smolen, J. S., Aletaha, D. & McInnes, I. B. Rheumatoid arthritis. Lancet 388, 2023–2038 (2016).

4. Rinn, J. L., Bondre, C., Gladstone, H. B., Brown, P. O. & Chang, H. Y. Anatomic Demarcation by Positional Variation in Fibroblast Gene Expression Programs. PLoS Genet 2, e119 (2006).

5. Frank-Bertoncelj, M. et al. Epigenetically-driven anatomical diversity of synovial fibroblasts guides joint-specific fibroblast functions. Nat Commun 8, 14852 (2017).

6. Müller-Ladner, U. et al. Synovial fibroblasts of patients with rheumatoid arthritis attach to and invade normal human cartilage when engrafted into SCID mice. Am. J. Pathol. 149, 1607–1615 (1996).

7. Croft, A. P. et al. Rheumatoid synovial fibroblasts differentiate into distinct subsets in the presence of cytokines and cartilage. Arthritis Res. Ther. 18, 270 (2016).

8. Gerlag, D. M., Norris, J. M. & Tak, P. P. Towards prevention of autoantibody-positive rheumatoid arthritis: from lifestyle modification to preventive treatment. Rheumatology 55, 607–614 (2016).

9. Pap, T., Müller-Ladner, U., Gay, R. E. & Gay, S. Fibroblast biology. Role of synovial fibroblasts in the pathogenesis of rheumatoid arthritis. Arthritis Res. 2, 361–367 (2000).

10. Ospelt, C. & Gay, S. The role of resident synovial cells in destructive arthritis. Best Pract Res Clin Rheumatol 22, 239–252 (2008).

11. McGettrick, H. M., Butler, L. M., Buckley, C. D., Rainger, G. E. & Nash, G. B. Tissue stroma as a regulator of leukocyte recruitment in inflammation. J. Leukoc. Biol. 91, 385–400 (2012).

12. Mizoguchi, F. et al. Functionally distinct disease-associated fibroblast subsets in rheumatoid arthritis. Nat Commun 9, 789 (2018).

13. Stephenson, W. et al. Single-cell RNA-seq of rheumatoid arthritis synovial tissue using low-cost microfluidic instrumentation. Nat Commun 9, 791 (2018).

14. Choi, I. Y. et al. Stromal cell markers are differentially expressed in the synovial tissue of patients with early arthritis. PLoS ONE 12, e0182751 (2017).

15. Filer, A. The fibroblast as a therapeutic target in rheumatoid arthritis. Curr Opin Pharmacol 13, 413–419 (2013).

16. Roberts, E. W. et al. Depletion of stromal cells expressing fibroblast activation protein-α from skeletal muscle and bone marrow results in cachexia and anemia. J. Exp. Med. 210, 1137–1151 (2013).

17. Krenn, V. et al. Synovitis score: discrimination between chronic low-grade and high-grade synovitis. Histopathology 49, 358–364 (2006).

18. Aletaha, D. et al. 2010 Rheumatoid arthritis classification criteria: An American College of Rheumatology/European League Against Rheumatism collaborative initiative. Arthritis Rheum. 62, 2569–2581 (2010).

19. Filer, A. et al. Identification of a transitional fibroblast function in very early rheumatoid arthritis. Ann. Rheum. Dis. 76, 2105–2112 (2017).

20. Kouskoff, V. et al. Organ-Specific Disease Provoked by Systemic Autoimmunity. Cell 87, 811–822 (1996).

21. Sinha, R. et al. Index Switching Causes ‘Spreading-Of-Signal’ Among Multiplexed Samples In Illumina HiSeq 4000 DNA Sequencing. (2017). doi:10.1101/125724

22. Butler, A., Hoffman, P., Smibert, P., Papalexi, E. & Satija, R. Integrating single-cell transcriptomic data across different conditions, technologies, and species. Nat. Biotechnol. 36, 411–420 (2018).

23. Tirosh, I. et al. Single-cell RNA-seq supports a developmental hierarchy in human oligodendroglioma. Nature 539, 309–313 (2016).

24. Ashburner, M. et al. Gene ontology: tool for the unification of biology. The Gene Ontology Consortium. Nat. Genet. 25, 25–29 (2000).

25. Dobin, A. et al. STAR: ultrafast universal RNA-seq aligner. Bioinformatics 29, 15–21 (2013).

26. Love, M. I., Huber, W. & Anders, S. Moderated estimation of fold change and dispersion for RNA-seq data with DESeq2. Genome Biol. 15, 550 (2014).

27. Ross, E. A. et al. Treatment of inflammatory arthritis via targeting of tristetraprolin, a master regulator of pro-inflammatory gene expression. Ann. Rheum. Dis. annrheumdis–2016–209424 (2016). doi:10.1136/annrheumdis-2016-209424

28. Wehmeyer, C. et al. Sclerostin inhibition promotes TNF-dependent inflammatory joint destruction. Sci Transl Med 8, 330ra35–330ra35 (2016).

